# A Comprehensive Analysis of the Phylogenetic Signal in Ramp Sequences in 211 Vertebrates

**DOI:** 10.1101/2020.03.03.975433

**Authors:** Lauren M. McKinnon, Justin B. Miller, Michael F. Whiting, John S.K. Kauwe, Perry G. Ridge

**Affiliations:** Department of Biology, Brigham Young University, Provo, UT 84602 USA; M.L. Bean Museum, Brigham Young University, Provo, UT 84602 USA

**Keywords:** Codon usage bias, phylogenetics, phylogenomics, ramp sequence

## Abstract

**Background:** Ramp sequences increase translational speed and accuracy when rare, slowly-translated codons are found at the beginnings of genes. Here, the results of the first analysis of ramp sequences in a phylogenetic construct are presented.

**Methods:** Ramp sequences were compared from 211 vertebrates (110 Mammalian and 101 non-mammalian). The presence and absence of ramp sequences was analyzed as a binary character in a parsimony and maximum likelihood framework. Additionally, ramp sequences were mapped to the Open Tree of Life taxonomy to determine the number of parallelisms and reversals that occurred, and these results were compared to what would be expected due to random chance. Lastly, aligned nucleotides in ramp sequences were compared to the rest of the sequence in order to examine possible differences in phylogenetic signal between these regions of the gene.

**Results:** Parsimony and maximum likelihood analyses of the presence/absence of ramp sequences recovered phylogenies that are highly congruent with established phylogenies. Additionally, the retention index of ramp sequences is significantly higher than would be expected due to random chance (p-value = 0). A chi-square analysis of completely orthologous ramp sequences resulted in a p-value of approximately zero as compared to random chance.

**Discussion:** Ramp sequences recover comparable phylogenies as other phylogenomic methods. Although not all ramp sequences appear to have a phylogenetic signal, more ramp sequences track speciation than expected by random chance. Therefore, ramp sequences may be used in conjunction with other phylogenomic approaches.

## I. Introduction

THE central dogma of biology states that DNA is transcribed into RNA, which is subsequently translated in sets of three consecutive nucleotides, called codons [5]. Since there are 61 possible codons (and three stop codons) which encode only 20 amino acids, there is redundancy in the genetic code. Although synonymous codons encode the same amino acid, recent research has shown that translational efficiency differs between synonymous codons [6-8]. These differences cause a change in translational speed, which affects gene expression [9], [10]. Ramp sequences consist of 30-50 infrequent or slowly-translated codons (i.e., codons translated by a relatively small proportion of tRNA molecules from the tRNA pool) at the 5’ end of many genes [11]. These sequences serve as a means to regulate gene expression by evenly spacing ribosomes along the mRNA transcript to reduce downstream ribosomal collisions [11], [13] and reduce mRNA secondary structure at translation initiation [12].

We recently developed an algorithm, ExtRamp, to identify ramp sequences [14]. Previously, ramp sequences were known and characterized in only a few model species. ExtRamp identifies ramp sequences by calculating the relative codon efficiency of each codon and then estimating ribosomal speed at each location in the gene by computing the average codon efficiency within the ribosomal window. If an outlier portion is present at the beginning of the gene, it is considered a ramp sequence. Using this algorithm, species in most domains of life were shown to have an average of 10% of genes with ramp sequences [14]. Given the widespread presence of ramp sequences in most domains of life and their role in regulating translation, this project investigates the hypothesis that the presence or absence of a ramp sequence in a gene may be used as a morphological genomic character that can be used to recover a phylogenetic signal.

Phylogenies are essential to understanding the biological world and allow biologists to analyze similarities and differences between closely related species [1]. They also provide an evolutionary context to better understand biological processes and patterns. Our knowledge of phylogenetic relationships increases in accuracy as more data are found and more species are discovered and sequenced, which allows us to have greater clarity in our knowledge of past speciation events. In order to analyze the ever-increasing amounts of phylogenetic data, many methods and frameworks of phylogenetic inference have been developed, each of which seeks to determine species relationships according to a set of assumptions of evolutionary processes. Maximum likelihood is a statistical method that incorporates a model of evolution (transition and transversion frequencies, nucleotide frequencies, evolutionary rates, etc.) to limit the effects of homoplasy [2]. Parsimony seeks to maximize homology in phylogenies by minimizing *ad hoc* hypotheses of homoplasy [3]. The success of any method of phylogenetic inference depends on the accurate phylogenetic signal of the characters used. Nearly all characters, including morphological and molecular, contain some amount of homoplasy [4]. As additional phylogenetically informative characters are identified and analyzed in tree reconstruction methods, the estimated phylogenetic tree increases in accuracy. Therefore, the potential of ramp sequences as a novel phylogenetic character is analyzed in order to determine if ramp sequences provide useful signal in phylogenomic analyses. The aim of this project is primarily to determine if the presence of ramp sequences in genes is phylogenetically conserved. This project additionally investigates the possibility that ramp sequences display a different phylogenetic signal than other portions of the sequence. The potential for using ramp sequences in future phylogenomic studies is then evaluated.

## II. METHODS

### A. Data Collection and Processing

Reference genomes were downloaded along with their corresponding General Feature Format (GFF3) files from the National Center for Biotechnology Information (NCBI) database [15-18] in August 2018 using their FTP site: ftp://ftp.ncbi.nlm.nih.gov/genomes/refseq/. One reference genome was included from 217 vertebrate species. The mammalian taxonomic group was analyzed (114 mammalian species), as well as their vertebrate outgroup (103 non-mammalian species). The congruence of the phylogenetic signal of ramp sequences within mammalian species and their vertebrate outgroup is then assessed. All coding sequences (CDS) data were extracted from the reference genomes. Any annotated exceptions were filtered from the dataset such as translational exceptions, unclassified transcription discrepancies, and suspected errors from the data. The analysis included all genes that contained an ortholog annotation according to NCBI ortholog annotations. This included 34,202 orthologs for Mammalia and 41,337 orthologs for non-mammalian vertebrates.

### B. Identifying Ramp Sequences

Ramp sequences were identified using ExtRamp (Fig. 1). The relative codon adaptiveness was calculated for each codon by using its frequency in the genome. The translation rate at each codon in the gene is then estimated using the mean translational efficiency of a window of codons. A nine-codon sliding window was used to approximate the span of a ribosome. A ramp sequence is identified where translational bottlenecks occur at the beginning of a gene. ExtRamp was run on each of the FASTA files (.fasta) for the reference genomes using the options to output the ramp sequence and the portion after the ramp sequence as described in the README file (https://github.com/ridgelab/ExtRamp) The exact command used is included in Supplementary Note 1.

**Fig. 1:**
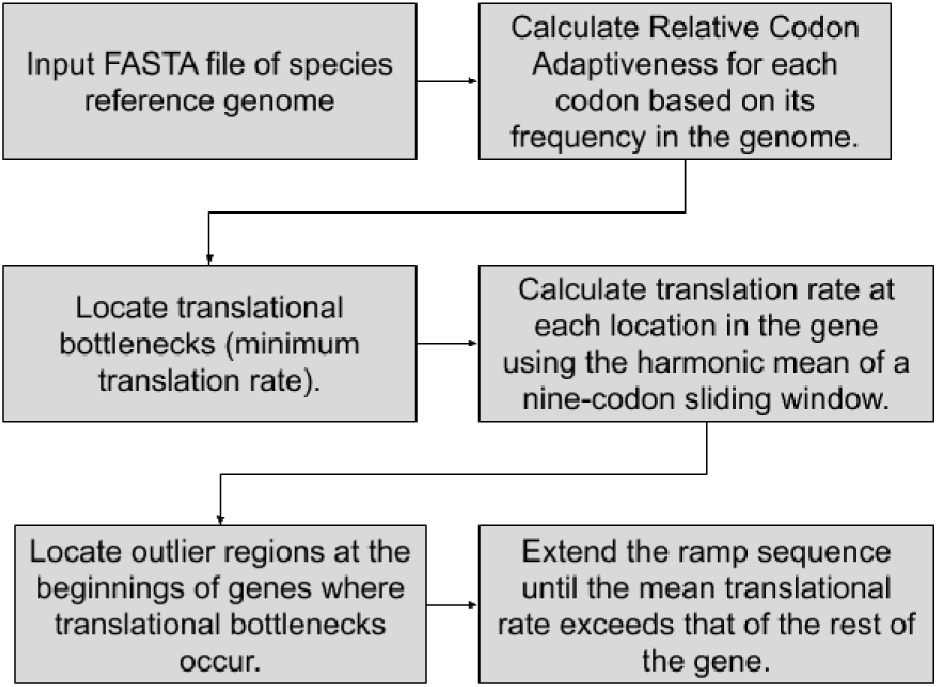
Flowchart for finding ramp sequences using ExtRamp.

### C. Recovering Phylogenies Using the Presence and Absence of Ramps

The presence of a ramp sequence was encoded in each identified ortholog in a binary matrix. If a ramp sequence was present in an ortholog, it was encoded in the matrix as a ‘1’, and if it was absent, it was encoded as a ‘0’. Any species that did not contain the ortholog was assigned a ‘?’ for a missing value. The effect of missing data was limited by applying a filter to the data. An orthologous group of ramp sequences was included in the analyses only if it was found in at least 5% of the species. A species was included in the analyses only if it contained at least 5% of the ramp sequences. After applying this filter, mammalian species had a mean of 16.31% ±7.81% and Non-mammalian vertebrates had a mean of 28.50% ± 13.11% missing data.

Phylogenetic trees were recovered in parsimony using Tree Analysis using New Technology (TNT) [19]. Most parsimonious trees were found by saving multiple trees using tree bisection reconnection (tbr) branch swapping [20]. Maximum likelihood trees were additionally recovered using IQTREE [21].

### D. Retrieving Reference Phylogenies

In order to determine the congruence of the phylogenetic signal of ramp sequences, each of the recovered phylogenies from Methods 2.3 were compared to established phylogenies from the Open Tree of Life (OTL) [22]. Although this phylogeny cannot be considered the “true” tree, it is created from a conglomeration of many phylogenetic studies, and thus provides a useful resource for benchmarking ramp sequences as a new character. Reference phylogenies were retrieved from the OTL using getOTLtree.py [23]. This program ref rences the OTL API to obtain OTL taxonomy identifiers for ach of the species which are used to retrieve the phylogeny from the OTL database. The exact command is included in Supplementary Note 2.

### E. Comparisons with Reference Phylogenies

The accuracy of the ramps phylogenies was assessed by comparing them to the OTL reference phylogeny. The difference was quantified using branch percent comparisons, as implemented by the Environment for Tree Exploration toolkit ete3 compare module [24], [25]. This metric computes the percentage of branch similarity between two trees, where a high percentage corresponds to more similar trees. This metric was selected because of its ability to compare large trees, including unrooted trees and trees with polytomies. The baseline performance of the ete3 branch percent identity metric was determined by comparing 1000 random permutations of the mammalian and other vertebrate topologies to the OTL.

### F. Scoring Ramp Sequences

Using the binary matrix of ramp sequences within each ortholog, the extent to which ramp sequences are homoplasious was quantified by mapping each ramp sequence to the OTL. For each ramp sequence, the species were divided into two partitions based on presence or absence of the ramp sequence. Since autapomorphies do not provide phylogenetic information, an orthologous ramp sequence was required to be present in at least two species and absent in at least two species, assuming a fully-resolved tree. For each ramp sequence, the number of parallelisms and reversals that occurred was quantified. Parallelisms occur when a character arises independently multiple times due to convergent evolution. Reversals occur when a derived character is lost or when the character reverts back to its ancestral state. A ramp sequence was determined to be orthologous if it separated species according to their relationships reported in the OTL, and if the total number of gain/loss events equaled one. The number of origin and loss events was then used to calculate the retention index for each ramp sequence [26], where a retention index of zero represents a fully homoplasious character, and a retention index of one represents a character in which none of the states are homoplasious.

### G. Comparison with Random Permutations

In order to determine how the retention index of the ramp sequences compare to what would be expected from random chance, random permutations were performed. The taxa in the OTL were shuffled 1000 times in order to generate random trees. The tree topology of the OTL was maintained in order to prevent any bias due to differences in tree structures. Then the retention indices of the ramp sequences were calculated for each random tree in order to create a null distribution of retention indices due to random chance. The actual mean retention index of the ramp was compared to this distribution in order to calculate an empirical p-value.

### H. Statistical Calculation of Completely Orthologous Ramps

For each orthologous ramp sequence, the probability was calculated that it would form a monophyletic group in agreement with the OTL topology by random chance. The species were divided into two partitions: those with ramp sequences, and those without ramp sequences. The probability was then calculated that the partitions would randomly divide into a monophyletic group according to the OTL topology using the method previously described in [27]. The probability that the first species will be assigned to a monophyletic group is 1. Then, the probability that the second species will be assigned to the monophyletic group is defined by the difference between the number of species in a taxonomic group and the number of species already assigned to that taxonomic group, which is then divided by the total number of species not yet assigned to a taxonomic group.

The probability is then calculated of assigning all species to the correct monophyletic group in (1), where s represents the number of species in the smaller partition and t represents the total number of species.

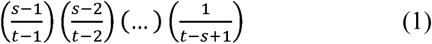

This equation can be simplified to (2).

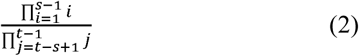

For example, if a ramp sequence contains three species in the smaller partition and five total species, the probability that this would form a monophyletic group in agreement with the OTL topology by random chance is the following:

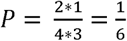

For each orthologous ramp sequence, the expected number of ramp sequences was calculated by multiplying the probability obtained from equation 3 by the total number of ramp sequences with that same taxonomic distribution (i.e. if the dataset contains 10 ramps where there are three species in the smaller partition and five total species, then the expected number is 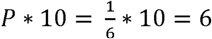) A chi square analysis was performed using the expected numbers versus the observed numbers in order to calculate a p-value.

### I. Recovering Phylogenies Using Aligned Sequence Data

In order to investigate the hypothesis that nucleotides in ramp sequences provide a different phylogenetic signal than other portions of the gene, the aligned sequences were analyzed using maximum likelihood and parsimony. Ramp sequences for each orthologous group were aligned using Clustal Omega [28]. Supplementary Note 3 contains the exact command.

A super-matrix was created by concatenating the aligned ramp sequences from each ortholog. If an ortholog was not present in a species, each nucleotide character for that sequence was encoded as a ‘?’ for missing data. The super-matrices were then used in IQ-TREE [29] to select the best model [30] and perform a Maximum Likelihood estimation of the phylogeny. The super-matrices were additionally used in TNT in order to recover phylogenies using parsimony.

Phylogenies were similarly recovered using the aligned sequence after the ramp and the complete gene sequence for each orthologous gene. For the maximum likelihood analysis, the size of the datasets for the portion after the ramp sequence and the complete sequence rendered the automatic model selection impractical due to computational demands. Therefore, the same models were selected that were used for the ramp analyses, which were GTR+F+R5 for Mammalia and GTR+F+R8 for non-mammalian vertebrates.

## III. RESULTS

### A. Data Analyzed

The original dataset consisted of a reference genome from each species in Mammalia and non-mammalian vertebrates from the NCBI database. A reference genome contains the most common sequences for each species. Ramp sequences were then extracted from all species. This dataset was filtered to include only ramp sequences that were identified in orthologous genes of at least 5% of the species. Also, a species was removed if it contained less than 5% of the included ramp sequences. Table I shows the resulting dataset that was included in the phylogenetic algorithms.

**TABLE I.** NUMBER OF SPECIES AND ORTHOLOGS ANALYZED

|  | Mammalia | Non-mammalian<br>Vertebrates |
| --- | --- | --- |
| **Species** | 114 | 133 |
| **Species containing ramp sequences** | 112 | 132 |
| **Species after filter** | 110 | 101 |
| **Orthologs** | 34,202 | 41,337 |
| **Orthologs containing ramp sequences** | 16,022 | 18,779 |
| **Orthologs after filter** | 11,670 | 8,450 |

The number of species and orthologs for each of the taxonomic groups are reported for the number of reference genomes contained in the NCBI database, the number that contained ramp sequences, and the number that passed the 5% filter criteria.

### B. Presence and Absence of Ramps Phylogenies

Phylogenies were recovered using each of the species and orthologs that passed the filter. A binary matrix was created where each ramp sequence was considered a character, with the character states of ramp present or absent. Species that did not contain an ortholog were coded as missing. Parsimony analyses were performed, and all maximum parsimony trees were retained. This resulted in two maximum parsimony trees for Mammalia, and two maximum parsimony trees for non-mammalian vertebrates (Supplementary Fig. 1 – 4). Each maximum parsimony tree was compared against the OTL taxonomy using the branch percent identity, and the resulting percentages were averaged for all of the maximum parsimony trees (Table II). Additionally, phylogenies were recovered using maximum likelihood (Supplementary Fig. 5 – 6). The branch percent identity is displayed in Table II. The table also provides branch percent comparisons of other tree reconstruction algorithms based on other genomic features to the OTL for reference as previously reported in [23]. These include Codon Aversion Motifs, Amino Acid Motifs, Codon Pairing, Feature Frequency Profiles, the word-based methods of CVTree, ACS, Andi, and Filter-spaced word matches, as well as maximum likelihood (Table II) [23], [29], [31]-[36]. A brief description of these algorithms is provided in Supplementary Table I.

**TABLE II.** BRANCH PERCENT IDENTITY COMPARED TO THE OTL

|  | Mammalia | Non-mammalian<br>vertebrates |
| --- | --- | --- |
| **Ramps – Parsimony** | 70 | 64 |
| **Ramps – Maximum likelihood** | 73 | 74 |
| **Codon Aversion Motifs** | 77 | 66 |
| **Amino Acid Motifs** | 63 | 56 |
| **Codon Pairing** | 90 | 77 |
| **Feature Frequency Profiles** | 52 | 54 |
| **CVTree** | 69 | 68 |
| **ACS** | 90 | 76 |
| **Andi** | 95 | 81 |
| **Filter-space word matches** | 94 | 80 |
| **Maximum likelihood** | 93 | 81 |

Branch percent identities of the presence/absence of ramps phylogenies compared to reference phylogenies.

The presence and absence of ramp sequences recovered 70 – 73% of relationships in the OTL for Mammalia, and 64 – 74% of relationships in the OTL for non-mammalian vertebrates. Ramp sequences perform comparably to the algorithms based on other genomic features reported.

### C. Retention Index of Ramp Sequences

The number of parallelisms and reversals for each ramp sequence was counted according to the OTL. This number was then used to calculate the retention index of each ramp sequence (Fig. 2).

**Fig. 2:**
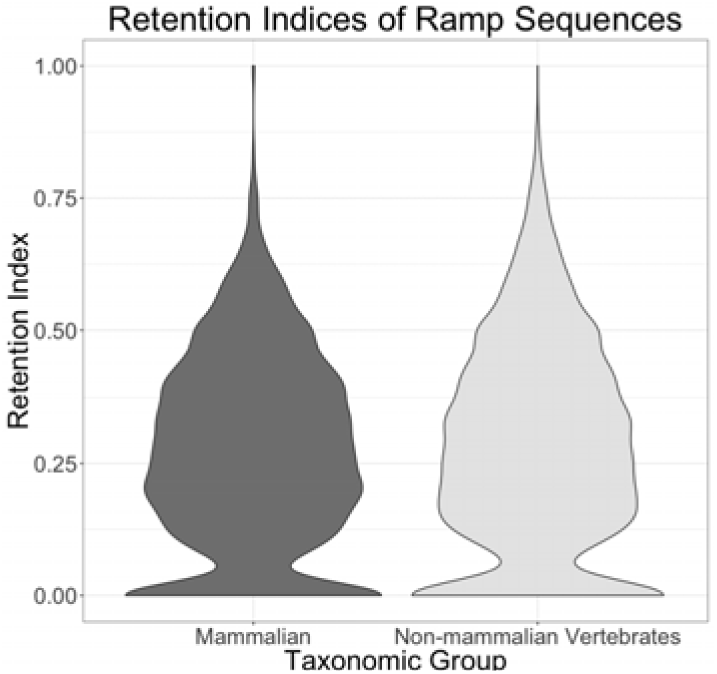
The retention index of each ramp sequence in non-mammalian vertebrates and Mammalia.

### D. Comparisons with Random Permutations

In order to determine how the retention index of ramp sequences compares to what would be expected by random chance, 1000 random permutations were performed by shuffling the species in the Open Tree of Life. The observed average retention index for Mammalia was higher than all the random permutations, for an empirical p-value of 0. This same result was observed for non-mammalian vertebrates (Fig. 3).

**Fig. 3:**
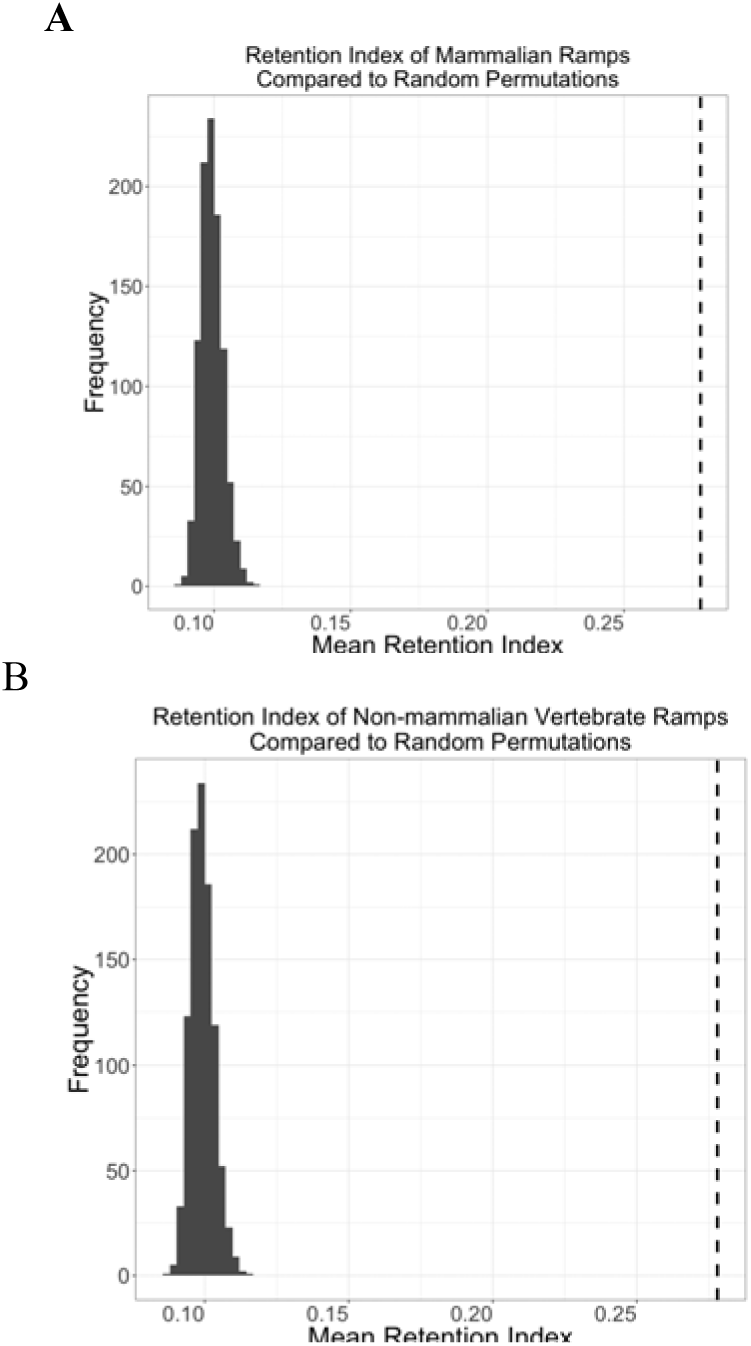
The mean retention index for 1000 random permutations in Mammalia (panel A) and Non-mammalian vertebrates (panel B). The actual mean retention index is represented with the dashed line.

### E. Orthologous Ramp Sequences and Statistical Probabilities

Orthologous ramp sequences were defined as all ramp sequences that had only one origin/loss event on the OTL. The statistical probability that each ramp sequence would be completely homologous by random chance was also calculated. The chi-square analysis of the observed number versus expected number of orthologous codons resulted in a p-value of zero for both Mammalia and non-mammalian vertebrates.

### F. Aligned Sequence Phylogenies

The filtered ramp sequences in orthologs were aligned and concatenated to make a super-matrix of nucleotide character data. This was done for the extracted ramp sequences, the portion after the ramp sequence, and the combined portions of the genes. Each of these super-matrices were used to recover phylogenetic trees using parsimony and maximum likelihood. These trees were then compared to the OTL taxonomy using the branch percent identities (Supplementary Fig. 7). Aligned ramp sequences showed lower congruence with the OTL than the portion after the ramp or the complete gene. However, the differences were not statistically significant.

## IV. DISCUSSION

Ramp sequences have been discovered relatively recently, and an algorithm for extracting ramp sequences was developed only last year. Given that an algorithm for extracting ramp sequences was only recently developed, the phylogenetic implications of ramp sequences have previously been unknown. The results of these analyses suggest that the presence or absence of a ramp sequence in some orthologs is largely congruent with the Open Tree of Life. However, some ramp sequences do not appear to be bound by the same evolutionary constraints. Therefore, while ramp sequences may add additional support to a phylogeny, they should be used in conjunction with other phylogenomic methods.

By considering only the presence or absence of a ramp sequence in orthologous genes in Mammalia and non-mammalian vertebrates, a parsimony analysis was able to recover a phylogeny that was relatively similar, as compared to other genomic approaches, to the OTL taxonomy for both Mammalia (70% to 73%) and non-mammalian vertebrates (64% to 74%). These results are comparable to those of Codon Aversion Motifs, Amino Acid Motifs, Feature Frequency Profiles, and CVTree. They are only slightly lower than Codon Pairing, ACS, Andi, Filter-spaced word matches and maximum likelihood. These results indicate that considering the presence or absence of a ramp sequence as a morphological genomic character is able to recover a comparable percentage of species relationships as other genomic features.

The analysis of parallelisms and reversals of ramp sequences suggest that ramp sequences contain phylogenetic information. According to the random permutations, the retention index is higher than would be expected due to random chance (p-value = 0) for both Mammalia and non-mammalian vertebrates. Additionally, both Mammalia and non-mammalian vertebrates had more completely orthologous ramp sequences as compared to the OTL than would be expected by random chance (p-value ≈ 0). This analysis shows that although some ramp sequences are homoplasious, the overall usage of ramp sequences shows statistically significant levels of homology as compared to the OTL topology.

Separate analyses were performed of aligned ramp sequences, portion after the ramp sequence, and complete sequence in order to compare the phylogenetic signal of different regions of genes. These analyses were completed using both maximum likelihood and parsimony, and the branch percent comparisons were compared for the OTL taxonomy. The results indicated that ramp sequences show less congruence to the OTL than the rest of the sequence portion. However, this result was not statistically significant and may have been confounded by the small length of the ramp sequence relative to the rest of the gene.

The analyses performed show that ramp sequence usage tracks speciation more frequently than random chance in Mammalia and non-mammalian vertebrates. Additionally, it is likely that ramp sequences will provide additional phylogenetic information for other taxonomic groups as more orthologs are identified and annotated across species. The results of these analyses show that ramp sequences may be used as a phylogenetic character state in future phylogenomic analyses.

## Acknowledgment

We appreciate contributions of Brigham Young University and the Fulton Supercomputing Laboratory at Brigham Young University for supporting our research.

## References

[1] Haszprunar, G., The Types of Homology and their Significance for Evolutionary Biology and Phylogenetics. Journal of Evolutionary Biology, 1992. 5(1): p. 13–24.

[2] Felsenstein, J., Evolutionary trees from DNA sequences: a maximum likelihood approach. J Mol Evol, 1981. 17(6): p. 368–76.

[3] Farris, J., The logical basis of phylogenetic analysis. 1983.

[4] Sanderson, M.J. and L. Hufford, Homoplasy : the recurrence of similarity in evolution. 1996, San Diego: Academic Press. xxv, 339 p.

[5] Crick, F., Central dogma of molecular biology. Nature, 1970. 227(5258): p. 561–3.

[6] Halder, B., A.K. Malakar, and S. Chakraborty, Nucleotide composition determines the role of translational efficiency in human genes. Bioinformation, 2017. 13(2): p. 46–53.

[7] Hanson, G. and J. Coller, Codon optimality, bias and usage in translation and mRNA decay. Nat Rev Mol Cell Biol, 2018. 19(1): p. 20–30.

[8] Liu, H., et al., Codon usage bias in 5’ terminal coding sequences reveals distinct enrichment of gene functions. Genomics, 2017. 109(5-6): p. 506–513.

[9] Cohen, E., Z. Zafrir, and T. Tuller, A code for transcription elongation speed. RNA Biol, 2018. 15(1): p. 81–94.

[10] Tarrant, D. and T. von der Haar, Synonymous codons, ribosome speed, and eukaryotic gene expression regulation. Cell Mol Life Sci, 2014. 71(21): p. 4195–206.

[11] Tuller, T., et al., An evolutionarily conserved mechanism for controlling the efficiency of protein translation. Cell, 2010. 141(2): p. 344–54.

[12] Goodman, D.B., G.M. Church, and S. Kosuri, Causes and effects of N-terminal codon bias in bacterial genes. Science, 2013. 342(6157): p. 475–9.

[13] Tuller, T. and H. Zur, Multiple roles of the coding sequence 5’ end in gene expression regulation. Nucleic Acids Res, 2015. 43(1): p. 13–28.

[14] Miller, J.B., L.R. Brase, and P.G. Ridge, ExtRamp: a novel algorithm for extracting the ramp sequence based on the tRNA adaptation index or relative codon adaptiveness. Nucleic Acids Res, 2019. 47(3): p. 1123–1131.

[15] Coordinators, N.R., Database resources of the National Center for Biotechnology Information. Nucleic Acids Res, 2013. 41(Database issue): p. D8–D20.

[16] Pruitt, K.D., et al., RefSeq: an update on mammalian reference sequences. Nucleic Acids Res, 2014. 42(Database issue): p. D756–63.

[17] Pruitt, K.D., et al., Introducing RefSeq and LocusLink: curated human genome resources at the NCBI. Trends Genet, 2000. 16(1): p. 44–7.

[18] Tatusova, T., et al., RefSeq microbial genomes database: new representation and annotation strategy. Nucleic Acids Res, 2014. 42(Database issue): p. D553–9.

[19] Goloboff, P.A., J.S. Farris, and K.C. Nixon, TNT: Tree Analysis Using New Technology. 2005. 54: p. 176–178.

[20] Kumar, S., et al., MEGA X: Molecular Evolutionary Genetics Analysis across Computing Platforms. Mol Biol Evol, 2018. 35(6): p. 1547–1549.

[21] Nguyen, L.T., et al., IQ-TREE: A Fast and Effective Stochastic Algorithm for Estimating Maximum-Likelihood Phylogenies. Molecular Biology and Evolution, 2015. 32(1): p. 268–274.

[22] Hinchliff, C.E., et al., Synthesis of phylogeny and taxonomy into a comprehensive tree of life. Proc Natl Acad Sci U S A, 2015. 112(41): p. 12764–9.

[23] Miller, J.B., et al., CAM: an alignment-free method to recover phylogenies using codon aversion motifs. PeerJ, 2019. 7: p. e6984.

[24] Huerta-Cepas, J., J. Dopazo, and T. Gabaldon, ETE: a python Environment for Tree Exploration. BMC Bioinformatics, 2010. 11: p. 24.

[25] Huerta-Cepas, J., F. Serra, and P. Bork, ETE 3: Reconstruction, Analysis, and Visualization of Phylogenomic Data. Mol Biol Evol, 2016. 33(6): p. 1635–8.

[26] Farris, J.S., The Retention Index and the Rescaled Consistency Index. Cladistics-the International Journal of the Willi Hennig Society, 1989. 5(4): p. 417–419.

[27] Miller, J.B., et al., Codon use and aversion is largely phylogenetically conserved across the tree of life. Mol Phylogenet Evol, 2020. 144: p. 106697.

[28] Sievers, F. and D.G. Higgins, Clustal Omega for making accurate alignments of many protein sequences. Protein Sci, 2018. 27(1): p. 135–145.

[29] Nguyen, L.T., et al., IQ-TREE: a fast and effective stochastic algorithm for estimating maximum-likelihood phylogenies. Mol Biol Evol, 2015. 32(1): p. 268–74.

[30] Posada, D. and K.A. Crandall, MODELTEST: testing the model of DNA substitution. Bioinformatics, 1998. 14(9): p. 817–8.

[31] Jun, S.R., et al., Whole-proteome phylogeny of prokaryotes by feature frequency profiles: An alignment-free method with optimal feature resolution. Proc Natl Acad Sci U S A, 2010. 107(1): p. 133–8.

[32] Miller JB, McKinnon LM, Whiting MF, Ridge PG. Codon Pairs are Phylogenetically Conserved: Codon pairing as a new class of phylogenetic characters. bioRxiv. 2019:654947.

[33] Zuo, G. and B. Hao, CVTree3 Web Server for Whole-genome-based and Alignment-free Prokaryotic Phylogeny and Taxonomy. Genomics Proteomics Bioinformatics, 2015. 13(5): p. 321–31.

[34] Ulitsky, I., et al., The average common substring approach to phylogenomic reconstruction. J Comput Biol, 2006. 13(2): p. 336–50.

[35] Haubold, B., F. Klotzl, and P. Pfaffelhuber, andi: fast and accurate estimation of evolutionary distances between closely related genomes. Bioinformatics, 2015. 31(8): p. 1169–75.

[36] Leimeister, C.A., S. Sohrabi-Jahromi, and B. Morgenstern, Fast and accurate phylogeny reconstruction using filtered spaced-word matches. Bioinformatics, 2017. 33(7): p. 971–979.

