## Supplementary Note 1 for "A Comprehensive Analysis of the Phylogenetic Signal in Ramp Sequences in 211 Vertebrates"

The following command generated a FASTA file of ramp sequences ${ramps_output} and a FASTA file of the sequence after the ramp ${after_output}.

python3 ExtRamp.py -i ${ref_fasta} -o ${ramps_output} -x ${after_output}
