## Supplementary Note 2 for "A Comprehensive Analysis of the Phylogenetic Signal in Ramp Sequences in 211 Vertebrates"

Reference phylogenies were retreived from the OTL database using the getOTL.py as found in the README file (<https://github.com/ridgelab/cam>), where ${input} is a list of species, and ${output} is the output file containing the phylogeny.

python getOTLtree.py -i ${input} -o ${output}
