## Supplementary figures and images for "A Comprehensive Analysis of the Phylogenetic Signal in Ramp Sequences in 211 Vertebrates"

### Supplementary Figure 1

***Supplementary Fig. 1:***

Mammals parsimony tree #1

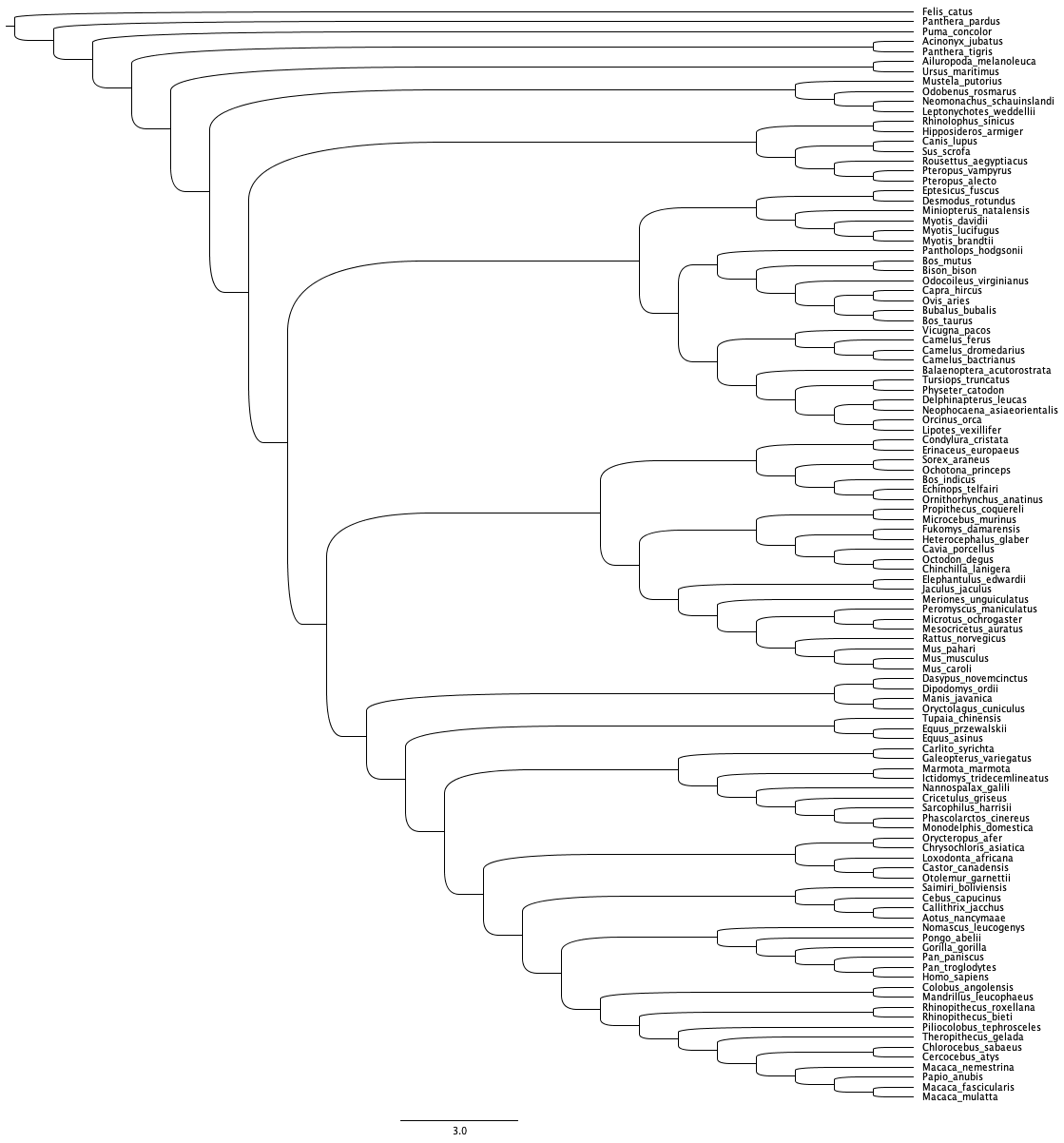

### Supplementary Figure 2

***Supplementary Fig. 2:***

Mammals parsimony tree #2

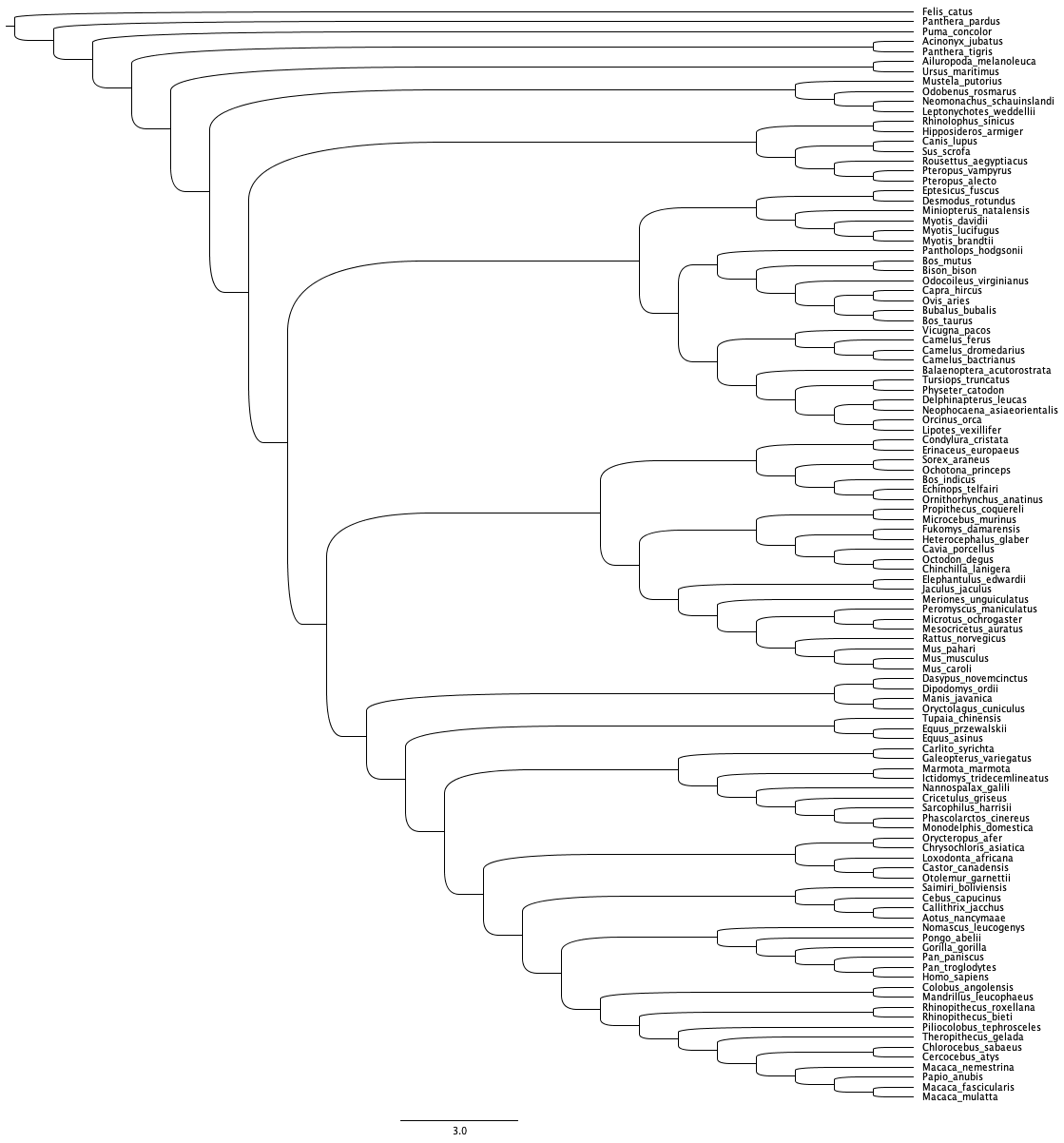

### Supplementary Figure 3

***Supplementary Fig. 3:***

Mammals maximum likelihood tree

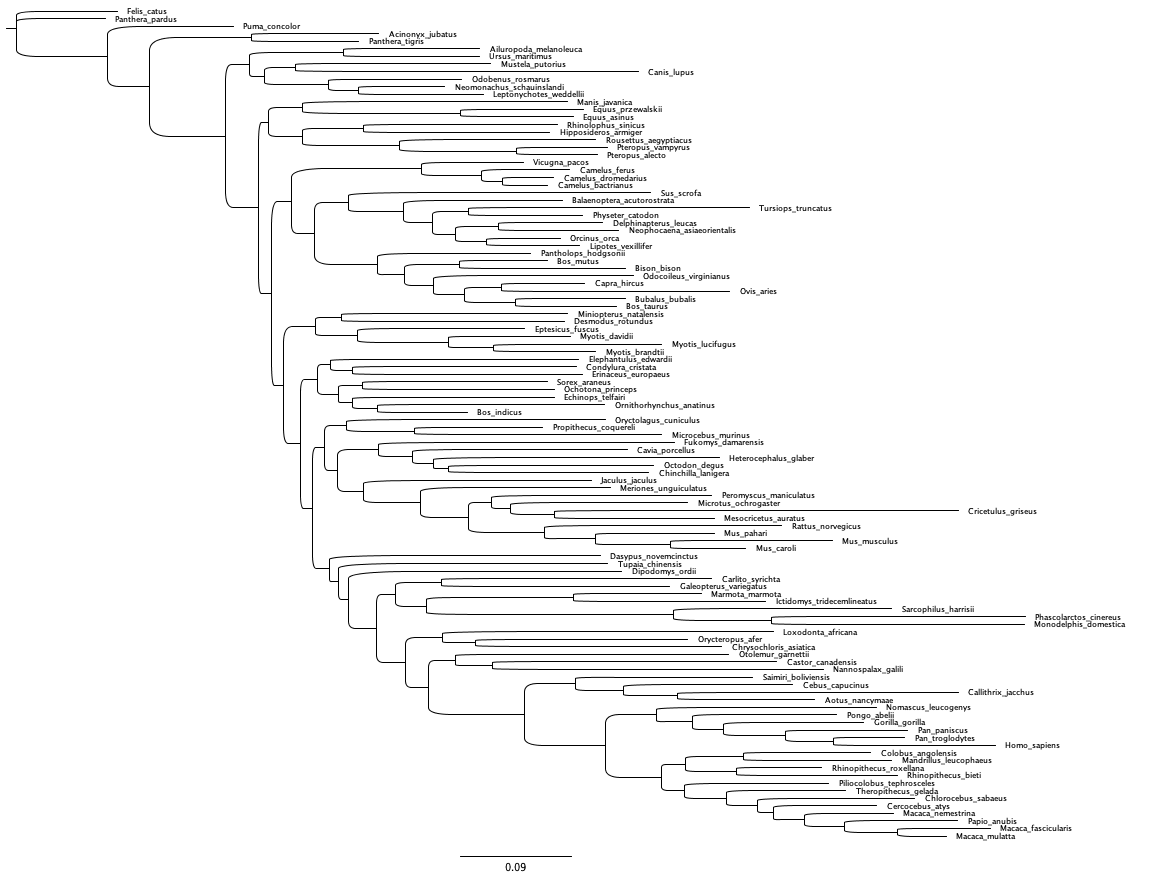

### Supplementary Figure 4

***Supplementary Fig. 4:***

Non-mammalian vertebrates parsimony tree 1
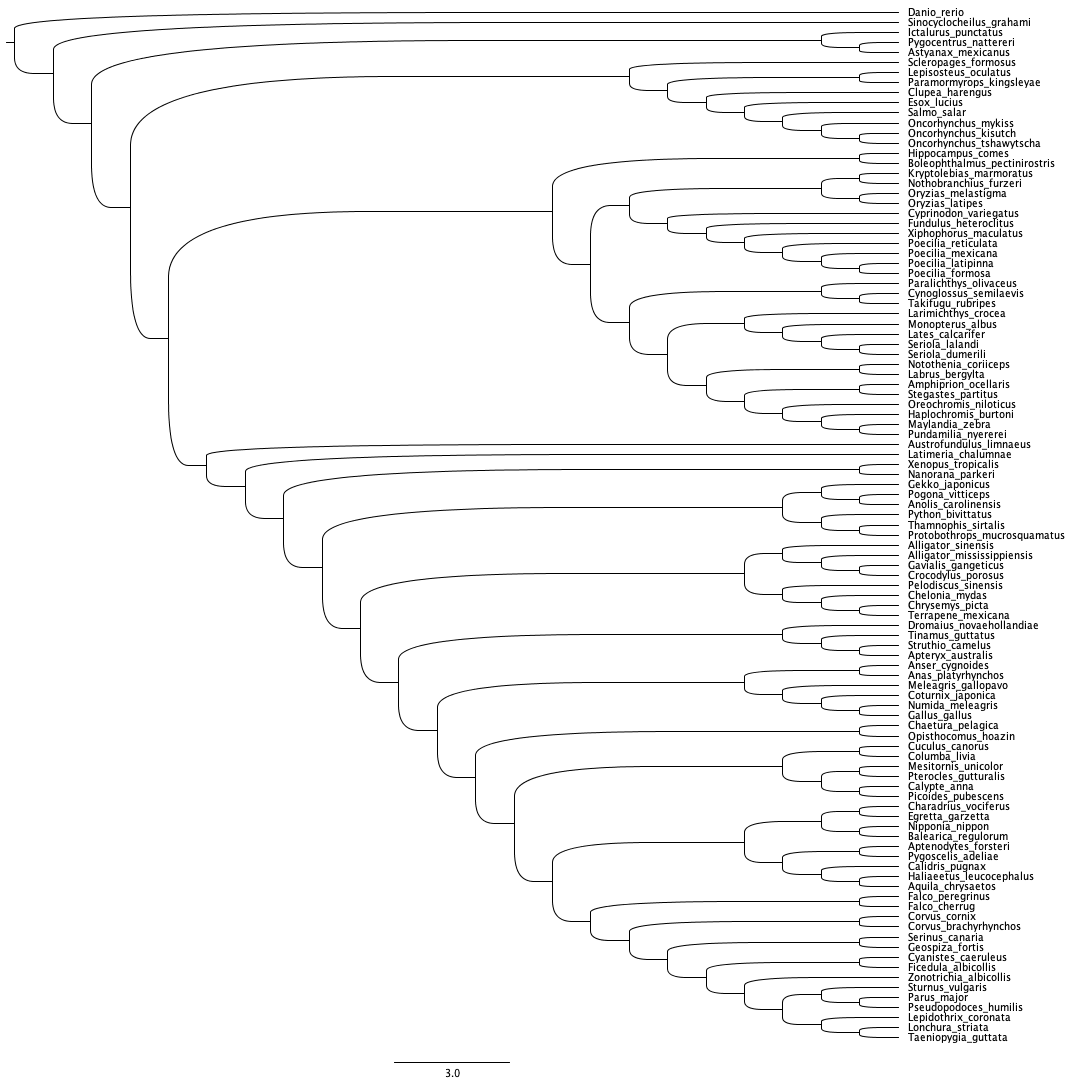

### Supplementary Figure 5

***Supplementary Fig. 5:***

Non-mammalian vertebrates parsimony tree 2

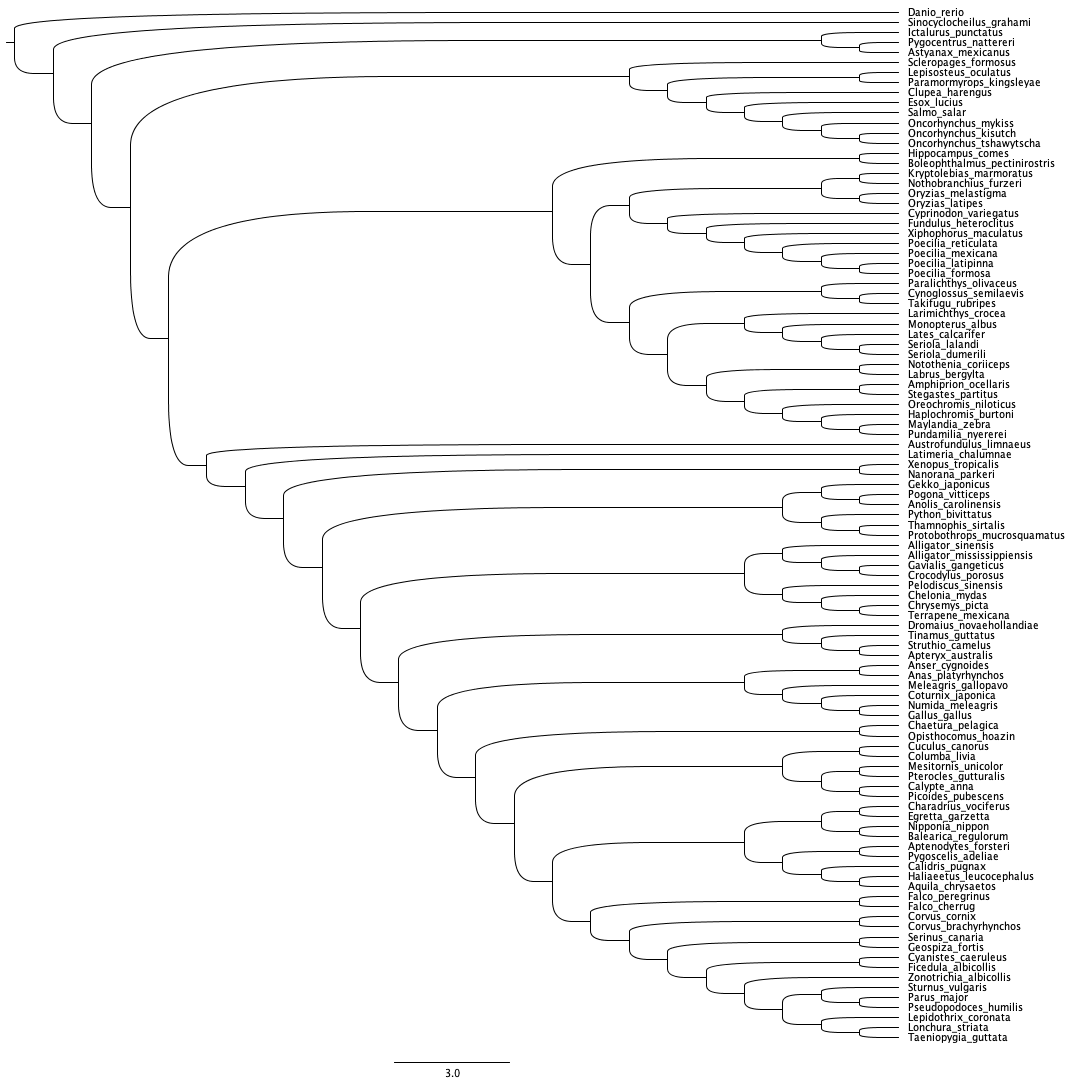

### Supplementary Figure 6

***Supplementary Fig. 6:***

Non-mammalian vertebrates maximum likelihood tree

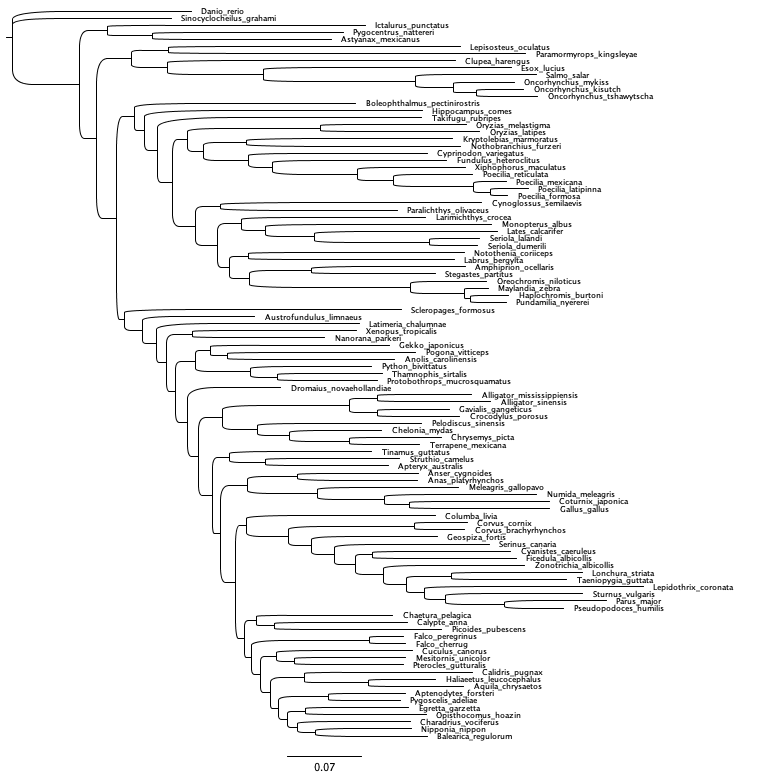
