## Supplementary Figure 7 for "A Comprehensive Analysis of the Phylogenetic Signal in Ramp Sequences in 211 Vertebrates"

***Supplementary Fig. 7:***

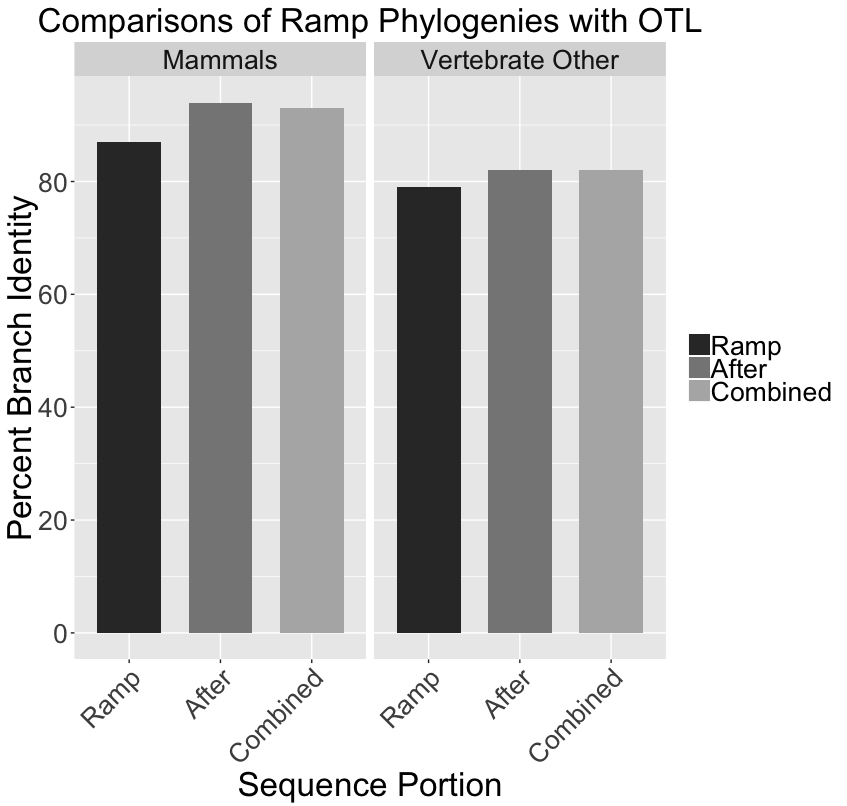

Phylogenies recovered using ramp sequences, the portion after the ramp sequence, and full sequences were compared to the OTL taxonomy using branch percent identity.
