## Supplementary Table 1 for "A Comprehensive Analysis of the Phylogenetic Signal in Ramp Sequences in 211 Vertebrates"

SUPPLEMENTARY TABLE I

BRIEF DESCRIPTION OF PHYLOGENETIC ALGORITHMS

| **Algorithm** | **Description** |
| --- | --- |
| Codon Aversion Motifs | Motifs of codons that are completely avoided are constructed for each gene. The distance between two species is calculated as the intersection of codon aversion motifs divided by the total number of unique motifs in the species with less motifs. |
| Amino Acid Motifs | Distance is calculated in the same manner as in Codon Aversion Motifs using amino acids instead of codons. |
| Codon Pairing | Codon pairing occurs when two codons that encode the same amino acid are located within a ribosomal window. Codon pairing was analyzed in a parsimony framework to infer a phylogeny. |
| Feature Frequency Profiles | The frequency of different k-mers is calculated, and the resulting profiles are compared between species to calculate a distance. |
| CVTree | Frequencies of words of a given length are calculated using composition vectors and then normalized based on the expected frequencies predicted by random chance. These frequencies are used to calculate a distance between species. |
| ACS | At each index of a gene, the longest matching substring is found in the second sequence. The average of these matching substrings is used to calculate a distance. |
| Andi | Andi creates micro-alignments between two sequences. It searches for mismatches that are bracketed by long, exact matches. These mismatches are then combined into a single matrix to estimate a mutation rate. |
| Filter-spaced word matches | Filter-spaced word matches finds matching-spaced words between sequences in a similar manner to Andi. It then adjusts the mutation rate, accounting for pattern matches caused by random chance. |
| Maximum Likelihood | This common alignment-based technique determines species relationships by finding the most likely phylogenetic tree by incorporating a model of evolution that includes estimated parameters such as transition/transversion frequencies, nucleotide frequencies, etc. |
